# Chemical identification of 18-hydroxycarlactonoic acid as an LjMAX1 product and conversion of 18-hydroxylcarlactonoates to canonical and non-canonical strigolactones in *Lotus japonicus*

**DOI:** 10.1101/837708

**Authors:** Narumi Mori, Aika Sado, Xiaonan Xie, Kaori Yoneyama, Kei Asami, Yoshiya Seto, Takahito Nomura, Shinjiro Yamaguchi, Koichi Yoneyama, Kohki Akiyama

**Affiliations:** Department of Applied Life Sciences, Graduate School of Life and Environmental Sciences, Osaka Prefecture University, Sakai, Osaka 599-8531, Japan; Department of Bioproductive Science, Graduate School of Agriculture, Utsunomiya University, Utsunomiya 321-8505, Japan; Center for Bioscience Research and Education, Utsunomiya University, Utsunomiya 321-8505, Japan; Graduate School of Agriculture, Ehime University, Matsuyama, Ehime 790-8566, Japan; PRESTO, Japan Science and Technology Agency, Kawaguchi, Saitama 332-0012, Japan; Department of Biomolecular Sciences, Graduate School of Life Sciences, Tohoku University, Sendai 980-8577, Japan; Department of Agricultural Chemistry, School of Agriculture, Meiji University, Kawasaki, Kanagawa, 214-8571, Japan; Institute for Chemical Research, Kyoto University, Uji, Kyoto, 611-0011, Japan

**Keywords:** *Lotus japonicus*, Fabaceae, Strigolactone, 5-Deoxystrigol, 18-Hydroxylcarlactonoic acid, Lotuslactone, Cytochrome P450 (CYP), *Lotus japonicus* MORE AXILLARY GROWTH 1 (MAX1)

## Abstract

Strigolactones (SLs) are a group of plant apocarotenoids that act as rhizosphere signaling molecules for both arbuscular mycorrhizal fungi and root parasitic plants. They also regulate plant architecture as phytohormones. The model legume *Lotus japonicus* produces canonical 5-deoxystrigol (5DS) and non-canonical lotuslactone (LL). The biosynthesis pathways of the two SLs remain elusive. In this study, we characterized the *L. japonicus MAX1* homolog, *LjMAX1*, found in the *Lotus japonicus* genome assembly build 2.5. The *L. japonicus max1 LORE1* insertion mutant was deficient in 5DS and LL production. A recombinant LjMAX1 protein expressed in yeast microsomes converted carlactone (CL) to 18-hydroxycarlactonoic acid (18-HO-CLA) via carlactonoic acid (CLA). Identity of 18-HO-CLA was confirmed by comparison of the methyl ester derivative of the MAX1 product with the chemically synthesized methyl 18-hydroycarlactonoate (18-HO-MeCLA) using LC-MS/MS. (11*R*)-CL was detected as an endogenous compound in the root of *L. japonicus.* ^13^C-labeled CL, CLA, and 18-HO-MeCLA were converted to [^13^C]-5DS and LL in plant feeding experiments using *L. japonicus* WT. These results showed that LjMAX1 is the crucial enzyme in the biosynthesis of *Lotus* SLs and that 18-hydroxylated carlactonoates are precursors for SL biosynthesis in *L. japonicus*.

## 1. Introduction

Strigolactones (SLs) are a group of apocarotenoid compounds identified as germination stimulants of root parasitic plants about 50 years ago (Cook et al., 1966). They were later demonstrated to be rhizosphere signaling molecules for arbuscular mycorrhizal symbiosis (Akiyama et al., 2005; Besserer et al., 2006) and to represent a new class of phytohormones regulating plant architecture, including shoot branching and several other aspects of shoot and root development (Gomez-Roldan et al., 2008; Kapulnik and Koltai, 2014; Sun et al., 2015; Umehara et al., 2008). To date, approximately 30 SLs have been characterized from root exudates of various plant species (Yoneyama et al., 2018b). Natural SLs are carotenoid-derived compounds consisting of an (*R*)-configurated methylbutenolide ring (D-ring) linked by an enol ether bridge to a less conserved second moiety (Al-Babili and Bouwmeester, 2015; Jia et al., 2018; Wang and Bouwmeester, 2018). SLs are classified into two groups, canonical and non-canonical SLs, according to their variable second moieties. Canonical SLs possess a C_19_ basic skeleton consisting of a tricyclic lactone (ABC ring) and the enol-ether–D ring. They are further divided into strigol- and orobanchol-type SLs, which have an α - and a β-oriented C ring, respectively. Non-canonical SLs contain the enol-ether–D ring moiety but lack the A, B, or C ring. The different SL molecules may all display different biological activities when acting as symbiotic and parasitic signals as well as endogenous plant hormones (Akiyama et al., 2010; Boyer et al., 2012; Zwanenburg & Pospíšil, 2013; Nomura et al., 2013a; Umehara et al., 2015; Mori et al., 2016; Xie et al., 2019).

Much progress has been made over the past decade about biosynthesis of SLs. The sequential action of DWARF27 (D27), a carotenoid isomerase, CAROTENOID CLEAVAGE DIOXYGENASE 7 (CCD7) and CAROTENOID CLEAVAGE DIOXYGENASE 8 (CCD8) converts β-carotene to carlactone (CL) (Alder et al., 2012; Flematti et al., 2016; Seto et al., 2014). *Arabidopsis thaliana* MAX1/CYP711A1 catalyzes three-step oxidations of the C19 methyl group of CL to form carlactonoic acid (CLA) (Abe et al., 2014). In rice, one of MAX1 homologs, Os900 acts as a CL oxidase to stereoselectively convert CL to an orobanchol-type SL, 4-deoxyorobanchol (4DO), and another MAX1 homolog of rice, Os1400, as orobanchol synthase, converts 4DO to orobanchol (Zhang et al., 2014). To determine which MAX1 reaction is conserved in the plant kingdom, the enzymatic function of MAX1 homolog in *Arabidopsis*, rice, maize, tomato, poplar and *Selaginella moellendorffii* were investigated (Kaori Yoneyama et al., 2018a; Zhang et al., 2018). The conversion of CL to CLA was found to be a common reaction catalyzed by MAX1 homolog. The ability of MAX1 homolog to metabolize CL can be classified into three types: A1-type, converting CL to CLA; A2-type, converting CL to 4DO via CLA; and A3-type, converting CL to CLA and 4DO to orobanchol. A new compound, most likely 18-hydroxycarlactonoic acid ((2*E*,3*E*)-4-(2-(hydroxymethyl)-6,6-dimethylcyclohex-1-en-1-yl)-2-((((*R*)-4-methyl-5-oxo-2,5-dihydrofuran-2-yl)oxy)methylene)but-3-enoic acid, 18-HO-CLA) was identified as a metabolite of CL by Os900, *Selaginella moellendorffii* MAX1 a/b and poplar MAX1 a/b. In the later stage of SL biosynthesis downstream of MAX1, *Arabidopsis* produces methyl carlactonoate (MeCLA) to CLA. MeCLA is further converted by LATERAL BRANCHING OXIDOREDUCTASE (LBO) enzyme, to an unidentified SL-like compounds (Brewer et al., 2016). LBO function may be involved to produce the major SL-type shoot branching hormone from MeCLA in the later steps of SL biosynthesis. In sorghum, a sulfotransferase, LOW GERMINATION STIMULANT 1 (LGS1), is likely to be involved in the formation of a strigol-type SL, 5-deoxystrigol (5DS) (Gobena et al., 2017).

It has been presumed that the biosynthesis of 5DS likely proceeds via the same mechanism with A2-type MAX1s, but is catalyzed by homologs that form the C ring with the β-configuration (Zhang et al., 2014; Al-Babili & Bouwmeester, 2015). However, a MAX1 homolog that converts CL to 5DS has not been discovered so far. It has remained elusive how strigol-type SLs are produced in plants. The conversion of CL and CLA to strigol-type SLs was shown by feeding experiments using sorghum, cotton and moonseed that are strigol-type SL producers (Iseki et al., 2018).

The model legume *Lotus japonicus* produces canonical 5DS that was isolated as the first hyphal branching factor for AM fungi (Akiyama et al., 2005). A non-canonical SL lotuslactone (LL) was recently identified as the second branching factor in *L. japonicus* (Xie et al., 2019). LL lacks the C-ring and has a seven-membered cycloheptadiene A-ring as in medicaol, a major SL of *Medicago truncatula* (Tokunaga et al., 2015). 5DS is strong inducer of hyphal branching in the AM fungus *Gigaspora margarita*, while LL is moderately active on the AM fungus. Among the basic SL biosynthesis pathway enzymes in *L. japonicus*, only a CCD7 homolog has been characterized (Liu et al., 2013). In *L. japonicus*, one MAX1 homolog (*LjMAX1*/CYP711A9) (Nelson, 2009) was found in the *Lotus japonicus* genome assembly build 2.5 (*Lotus japonicus* genome sequence project: http://www.kazusa.or.jp/lotus/) (Challis et al., 2013), but the function of the enzyme encoded by *LjMAX1* has not yet been characterized. Considering the biochemical function of MAX1 homologs of various plants (Abe et al., 2014; Kaori Yoneyama et al., 2018a; Zhang et al., 2014, 2018), we hypothesized that LjMAX1 is required for the oxidation of the SL precursor CL. In this study, to elucidate the enzymatic function of LjMAX1 in SL biosynthesis, we performed *in vitro* conversion of CL and CLA using a recombinant protein expressed in yeast microsomes. To confirm putative 18-HO-CLA that was detected as an LjMAX1 product, we chemically synthesized 18-HO-MeCLA and identified it by LC-MS/MS analysis. We then conducted feeding experiments with ^13^C-labeled CL, CLA and 18-HO-MeCLA to check whether these chemicals are precursors for 5DS and LL in *L. japonicus*.

## 2. Results

### 2.1. The cloning and characterization of LjMAX1

To confirm the involvement of *Lotus japonicus MAX1* homolog (chr1.CM0133.560.r2.m in build 2.5 from the ecotype Miyakojima MG-20) in the biosynthesis of SLs, one *LORE1* insertion mutants were obtained from the *Lotus* Base mutant collection (Fukai et al., 2012; Urbański et al., 2012; Małolepszy et al., 2016). In *ljmax1* (line 30101744), the *LORE1* retroelement was inserted in the first exon, 340 bp after the *LjMAX1* start codon, leading to very low accumulation of *LjMAX1* transcripts (Fig. S1 a,b). We analyzed the SL content in both WT and the *Ljmax1* mutant. We grew both genotypes in hydroponic solution without Pi to induce SL production. 5DS and LL were detected in the root exudates of WT, but were not detected in those of the *Ljmax1* mutant (Fig. S2). These results demonstrated that LjMAX1 is the crucial enzyme in the biosynthesis of *Lotus* SLs.

To characterize the enzymatic function of the *LjMAX1* in SL biosynthesis, we isolated the gene from *L. japonicus* ecotype Gifu B-129. The corresponding full length open reading frame was cloned by RT-PCR using total RNA prepared from seedling roots grown under Pi starvation. The open reading frame was 1617 bp encoding for a 539-amino acid protein. The amino acid sequence displayed 73% identity to Arabidopsis MAX1 (Fig. S3). It was discovered that the LjMAX1 gene in the Gifu B-129 ecotype of *L. japonicus* contains a single-base pair substitution of cytosine to thymine at position 1119 when compared with the gene in the Miyakojima MG-20 ecotype (Fig. S4). This is a synonymous substitution leaving unaffected the identity of the corresponding amino acid residue glycine in position 373.

### 2.2. *Lotus* MAX1 catalyzes the conversion of CL via CLA to 18-HO-CLA, but not to 5DS *in vitro*

To characterize the conversion of CL by LjMAX1, the recombinant protein was expressed in yeast strain WAT11 which was engineered to co-produce ATR1 using the galactose-inducible *GAL10-CYC1* promoter of yeast (Pompon et al., 1996). The resulting microsomal fractions were incubated with either CL or CLA, and their metabolites were analyzed by LC-MS/MS. When incubated with CL, LjMAX1 produced significant levels of CLA but not 5DS (Fig. 1). Although 5DS was not detected as a metabolite of CLA, the CLA+16 Da putatively identified as 18-HO-CLA was detected at small levels (Fig. 2) (Yoneyama et al., 2018a). To confirm the inability of LjMAX1 to produce 5DS, we carried out feeding experiments with CL or CLA by in vivo bioconversion using living yeast cells (Nomura et al., 2013b; Yoneyama et al., 2018a). Both substrates did not yield the product 5DS (Fig. S5a,b).

**Fig. 1.**
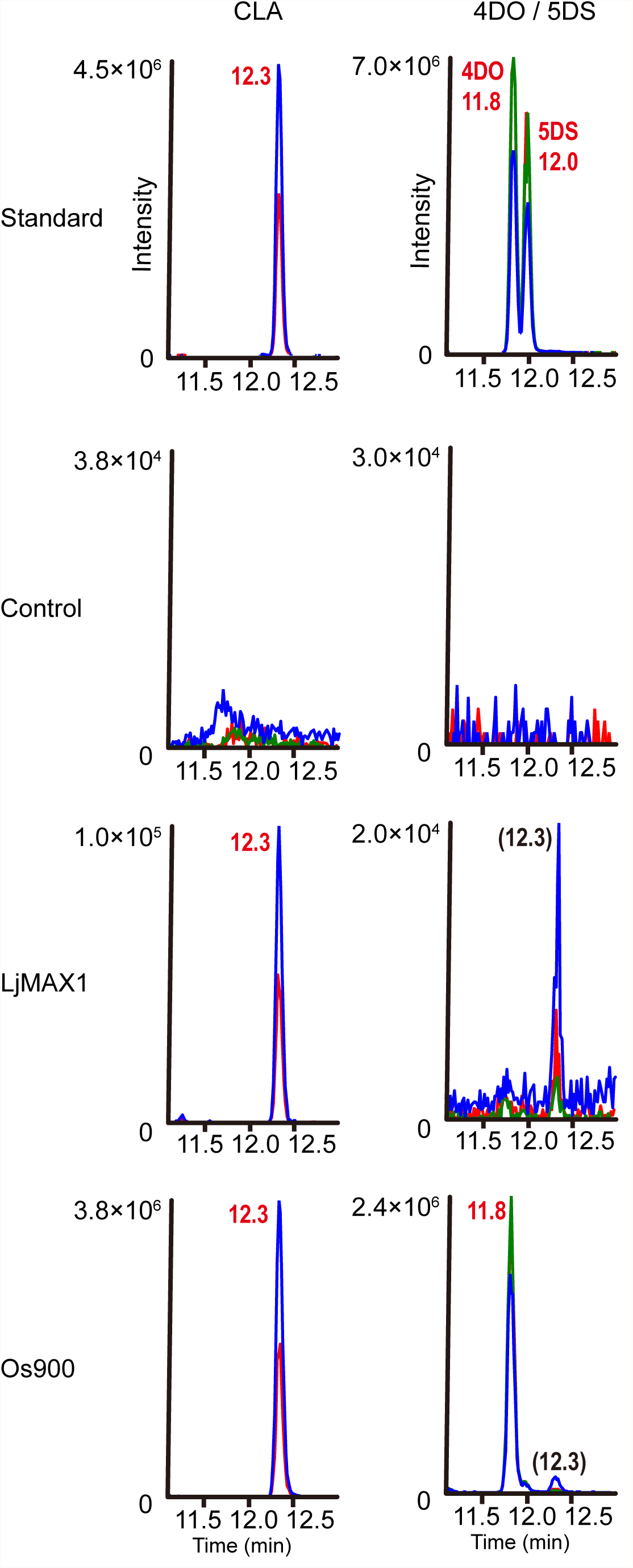
Detection of carlactonoic acid (CLA), 4-deoxyorobanchol (4DO) and 5-deoxystrigol (5DS) in carlactone (CL) feeding experiment by recombinant *Lotus japonicus* MAX1 (LjMAX1) and Os900. CL was incubated with yeast microsomes. Yeast microsomes having an empty vector and the expression vector pYeDP60-Os900 were used as a negative and a positive control, respectively. The extracts of the microsomes and authentic standard were analyzed by LC-MS/MS. MRM chromatograms of CLA (red: 331.10/69.00, blue: 331.10/113.00, *m/z* in negative mode), 4DO and 5DS (red: 331.15/216.00, blue: 331.15/97.00, green: 331.15/234.00, *m/z* in positive mode) are shown.

**Fig. 2.**
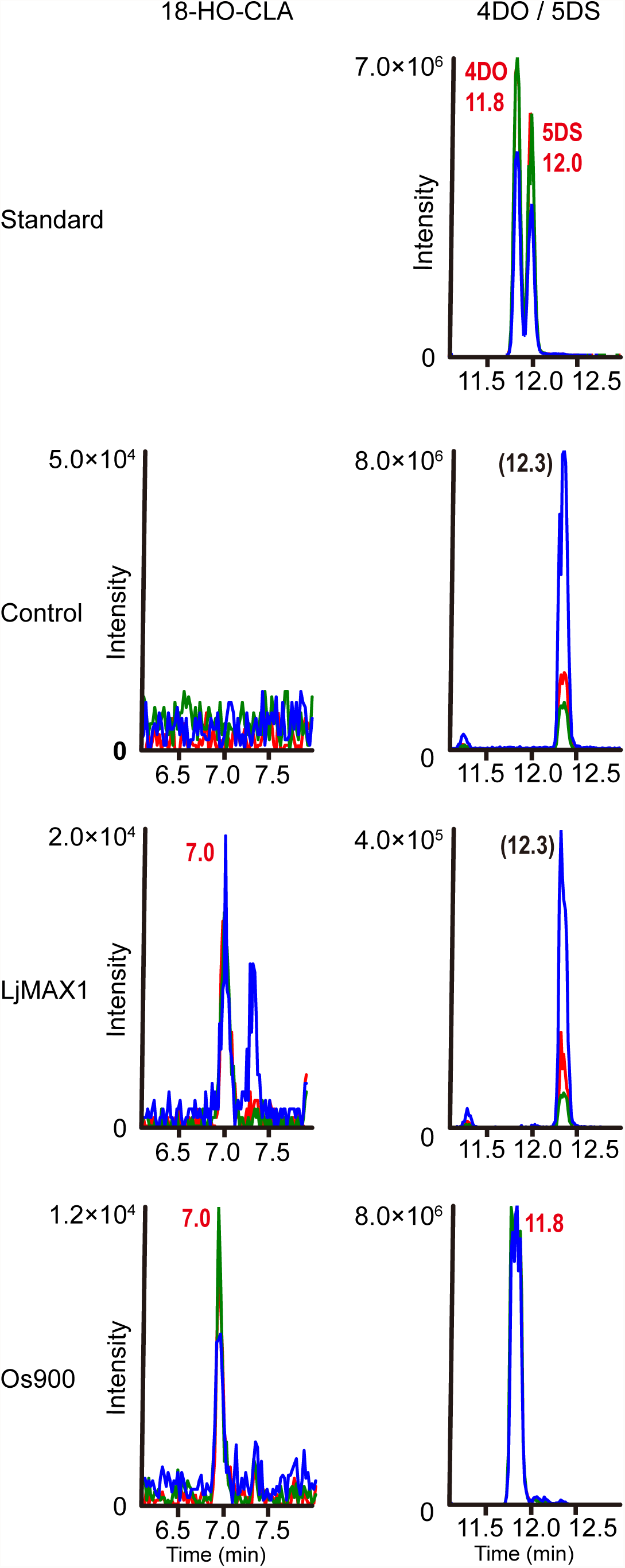
Detection of 18-hydroxycarlactonoic acid (18-HO-CLA), 4DO and 5DS in CLA feeding experiment by recombinant LjMAX1 and Os900. CLA was incubated with recombinant yeast microsomes. Yeast microsomes having an empty vector and the expression vector pYeDP60-Os900 were used as a negative and a positive control, respectively. The extracts of the microsomes and authentic standard were analyzed by LC-MS/MS. The peak at 7.1 min was presumed to be 18-HO-CLA which may be converted artificially to 4DO and 5DS in the ion source of mass spectrometer, and detected in the MRM transitions for 4DO and 5DS (red: 331.15/216.00, blue: 331.15/97.00, green: 331.15/234.00, *m/z* in positive mode).

### 2.3. Identification of the CLA+16 Da as 18-HO-CLA

To confirm the putative 18-HO-CLA, we synthesized 18-HO-MeCLA in eight steps (Scheme 1) starting from the triflate ester which was reduced with diisobutylaluminium hydride to give triflate alcohol. After protection of the hydroxyl group by a *tert*-butyl(dimethyl)silyl (TBDMS) group, the resultant silyl-protected triflate was subjected to the Heck reaction with acrolein to afford TBDMSO-C_12_-aldehyde. The silylated aldehyde was converted to TBDMSO-C_13_-aldehyde via the Corey-Chaykovsky epoxidation with dimethylsulfonium methylide, followed by methylaluminium bis(4-bromo-2,6-di-*tert*-butylphenoxide) (MABR)-promoted rearrangement of epoxide to aldehyde. The silylated C_13_-aldehyde was converted to the corresponding methyl ester with sodium cyanide and manganese oxide in methanol. Ester condensation of methyl ester with methyl formate followed by *O*-alkylation with bromo-butenolide provided 18-TBDMSO-MeCLA. After purification by preparative HPLC, 18-TBDMSO-MeCLA was deprotected with aqueous acetic acid to afford 18-HO-MeCLA. The identity of 18-HO-CLA was confirmed by comparison of retention time and mass spectra of the methyl ester derivative of the Os900 product with those of the chemically synthesized standard in LC-MS/MS analysis (Fig. 3).

**Fig. 3.**
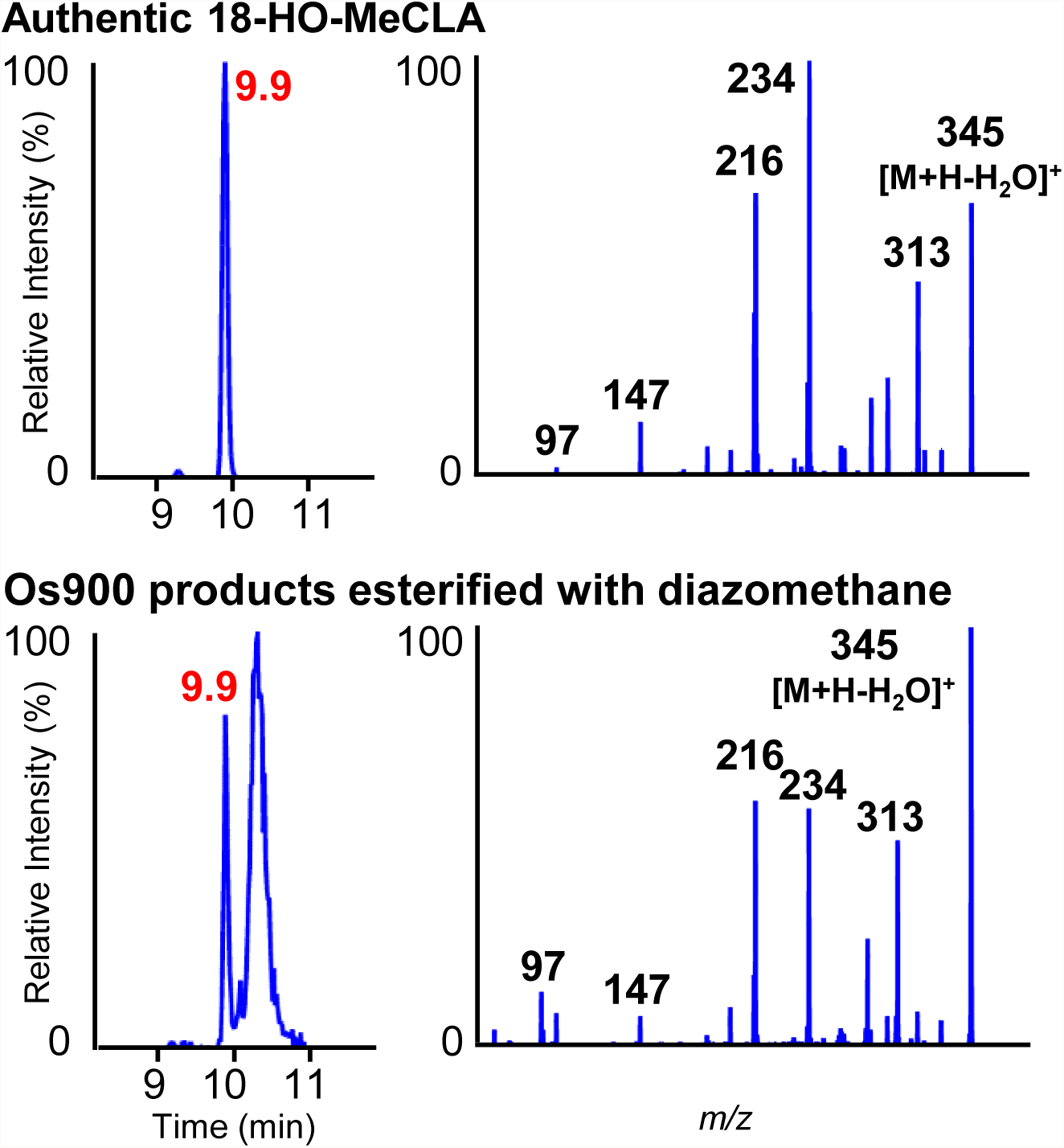
Identification of 18-HO-CLA as a MAX1 product. The CL metabolite by Os900 was methyl esterified with diazomethane and analyzed by LC-MS/MS. The product ions and retention time of the compound were identical to those of the authentic chemically synthesized methyl 18-hydroxycarlactonoate (18-HO-MeCLA). 18-HO-MeCLA was detected in the MRM transition 345.15[M+H-H_2_O]^+^/97.00, *m/z* in positive mode.

### 2.4. (11*R*)-CL is an endogenous precursor for (+)-5DS in *L. japonicus*

We analyzed endogenous CL in the root extract of *L. japonicus* WT using chiral LC-MS/MS. As in rice and *Arabidopsis*, only the (11*R*)-isomer of CL, but not the (11*S*)-isomer, was detected (Fig. S6). Next, to demonstrate that (+)-5DS was produced from (11*R*)-CL, we performed feeding experiments using *L. japonicus* WT while inhibiting endogenous SL production with the carotenoid pathway inhibitor fluridone (Matusova et al., 2005). The WT seedlings were grown hydroponically, and incubated with (11*R*)-or (11*S*)-[1-^13^CH_3_]-CL in the presence of fluridone. Significant incorporation of ^13^C-label into 5DS was observed for the (11*R*)-isomer but not for the (11*S*)-isomer (Fig. 4a). The absolute configuration of the resulting [8-^13^CH_3_]-5DS was determined to be (+)-5DS by comparison of the retention time with synthetic stereoisomers using chiral LC-MS/MS (Fig. 4b). These results showed that *L. japonicus* produced (+)-5DS from endogenous (11*R*)-CL.

**Fig. 4.**
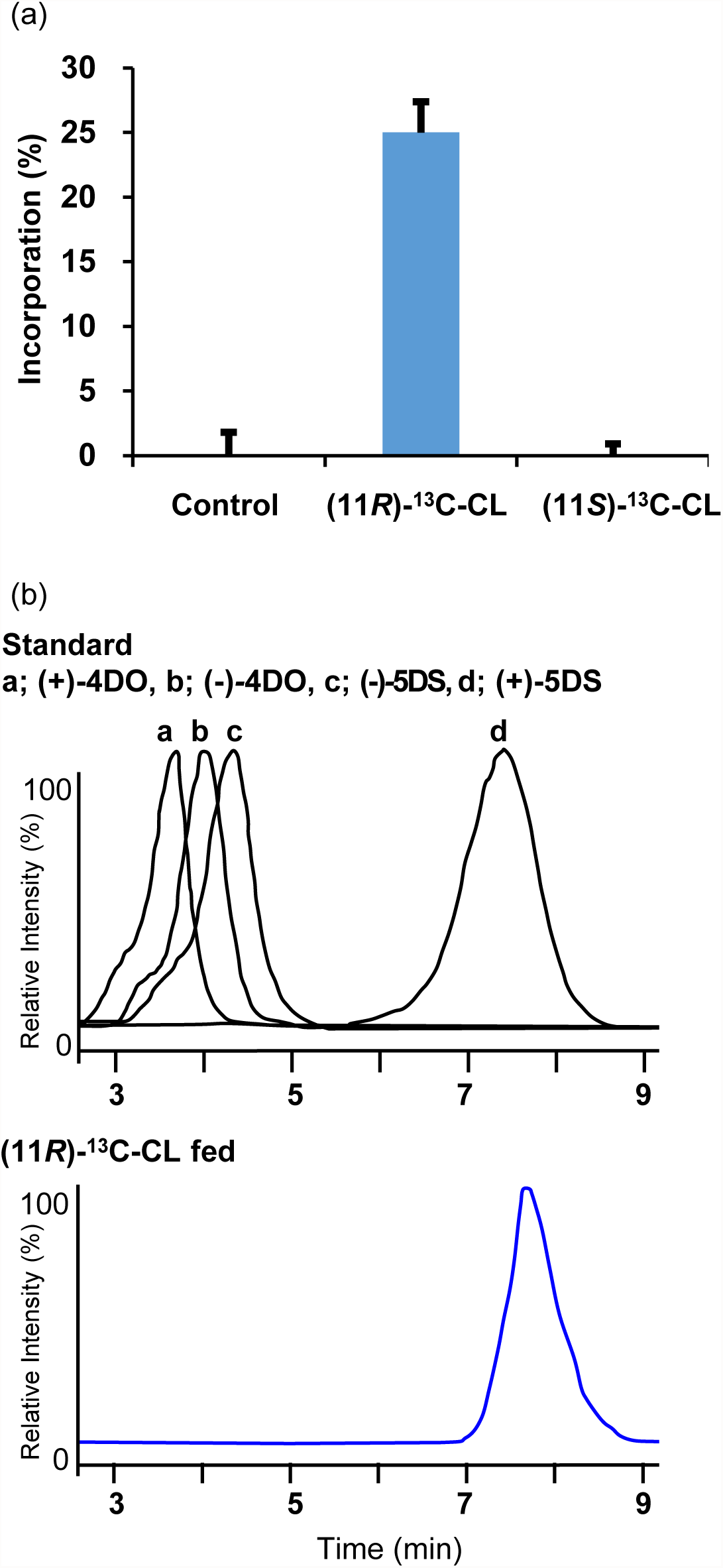
Conversion of [^13^C]-CL to [^13^C]-5DS in the feeding experiment using *L. japonicus* roots. (a) ^13^C-incorporation rates of (11*R*)- and (11*S*)-[1-^13^CH_3_]-CL into [8-^13^CH_3_]-5DS. Control was added acetone without substrates. ^13^C-incorporation rates were calculated from the formula: (peak area of *m/z* 332.15/97.00-peak area of *m/z* 331.15/97.00*0.20)/(peak area of 332.15/97.00+peak area of *m/z* 331.15/97.00), where 0.20 is the ratio of peak area of 332.15/97.00 to peak area of *m/z* 331.15/97.00 of natural 5DS. Peak areas were measured using the MRM transition for 5DS (331.15/97.00 and 332.15/97.00 *m/z* in positive mode) in LC-MS/MS analysis. *n* = 3. (b) Determination of the stereochemistry of 5DS converted from (11*R*)-[1-^13^CH_3_]-CL. Chiral LC-MS/MS analysis of stereoisomers of 5DS in root exudates after feeding of (11*R*)-[1-^13^CH_3_]-CL. MRM of synthetic standards of 5DS and 4DO stereoisomers and [^13^C]-(+)-5DS from *L. japonicus* root exudates after feeding (11*R*)-[1-^13^CH_3_]-CL.

### 2.5. CLA is converted to 5DS and 18-HO-CLA in *L. japonicus*

To investigate whether the LjMAX1 product CLA is a biosynthetic precursor for SLs *in planta*, we examined the conversion from exogenous CLA into SLs using *L. japonicus* WT. Feeding [1-^13^CH_3_]-CLA to fluridone-treated hydroponic culture was carried out, and labeled products were analyzed by LC-MS/MS. We could detect the peak of [8-^13^CH_3_]-5DS whose *m/z* is increased by one mass unit in [1-^13^CH_3_]-CLA-fed samples compared with an unlabeled authentic standard (Fig. 5a). In addition, [1-13CH_3_]-18-HO-CLA was detected as a metabolite of labeled CLA in root exudates based on the comparison of the full-scan MS spectrum and the retention time on LC with those of unlabeled authentic standard (Fig. 5b). In an attempt to detect 18-HO-CLA produced by *L. japonicus*, we could detect 18-HO-CLA by LC-MS/MS in the root exudates of WT plants, although the full-scan MS spectrum was not obtained due to its scarcity (Fig. S7). These results indicated that *L. japonicus* converts CLA to 5DS and 18-HO-CLA *in planta*.

**Fig. 5.**
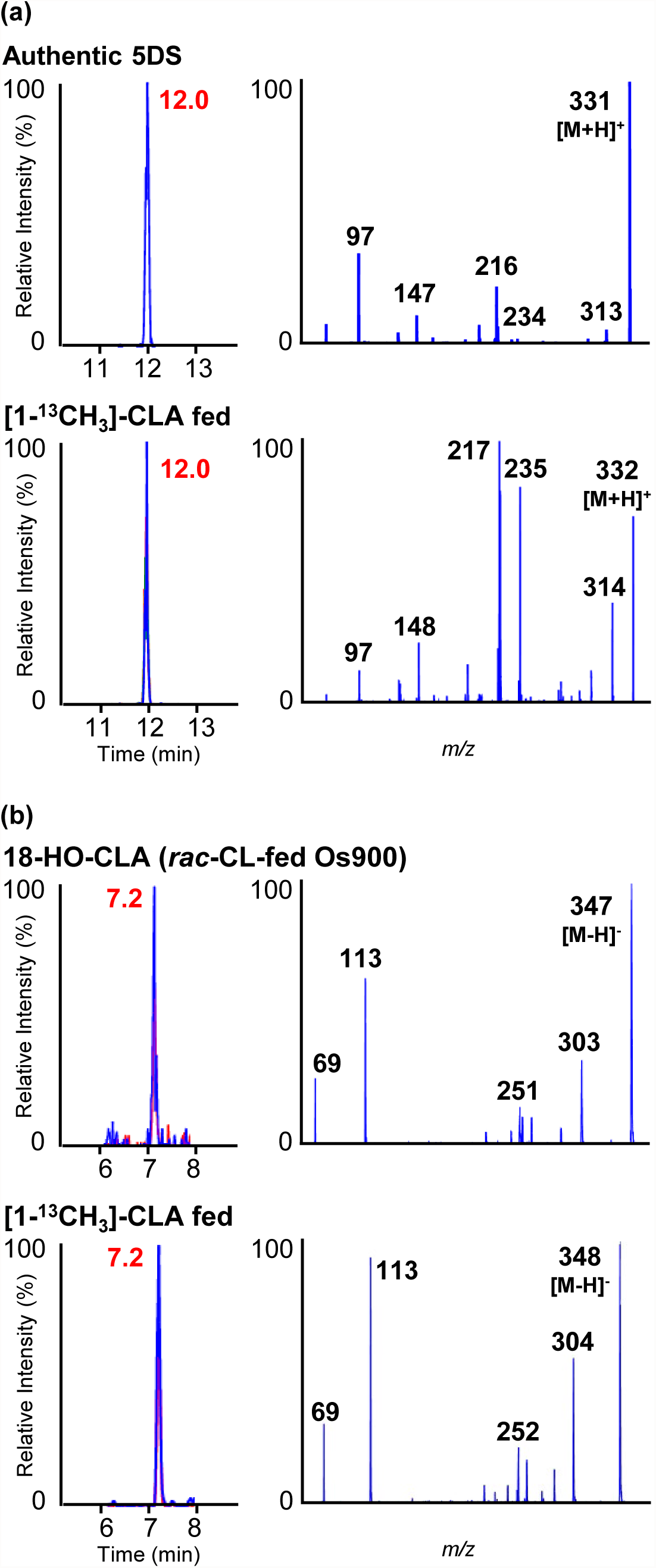
Conversion of [^13^C]-CLA to [^13^C]-5DS and [^13^C]-18-HO-CLA in the feeding experiment using *L. japonicus* roots. (a) Conversion of [1-^13^CH_3_]-CLA to [8-^13^CH_3_]-5DS. LC-MS/MS analysis of [8-^13^CH_3_]-5DS in root exudates after feeding [1-^13^CH_3_]-CLA. MRM chromatograms (left) and full-scan spectra of fragment ions (right). The MRM chromatograms of authentic 5DS (red: 331.15/217.00, blue: 331.15/97.00, green: 331.15/234.00, *m/z* in positive mode) and [8-^13^CH_3_]-5DS (red: 332.15/218.00, blue: 332.15/97.00, green: 332.15/235.00, *m/z* in positive mode) are shown. (b) Conversion of [1-^13^CH_3_]-CLA to [1-^13^CH_3_]-18-HO-CLA. LC-MS/MS analysis of [1-^13^CH_3_]-18-HO-CLA in root exudates after feeding [1-^13^CH_3_]-CLA. MRM chromatograms (left) and full-scan spectra of fragment ions (right). Authentic 18-HO-CLA was prepared by feeding CL to recombinant Os900. MRM chromatograms of authentic 18-HO-CLA (red: 347.00/303.00, blue: 347.00/113.00, *m/z* in negative mode) and [1-^13^CH_3_]-18-HO-CLA (red: 348.00/304.00, blue: 348.00/113.00, *m/z* in negative mode) are shown.

### 2.6. 18-HO-CLA derivatives are converted to 5DS and LL

We next conducted a feeding experiment to investigate whether the *in vitro* and *in planta* metabolite of CLA, 18-HO-CLA, is converted to 5DS in *L. japonicas*. Although 18-HO-CLA has been predicted to be an intermediate for canonical SLs, its chemical synthesis has not been achieved to date (Ueno et al., 2018; Kaori Yoneyama et al., 2018a; Zhang et al., 2014). Instead we synthesized [10-^13^C]-18-HO-MeCLA by the above established method using methyl [^13^C_1_]formate as a labeling reagent. We expected that methyl ester could be hydrolyzed to the corresponding carboxylic acid by non-specific esterases when fed to *L. japonicus* roots. A significant level of [10-^13^C]-18-HO-CLA was indeed detected in the root exudates of the WT plants, when [10-^13^C]-18-HO-MeCLA was fed to fluridone-treated hydroponic culture. Simultaneously, we could successfully detect [6 -^13^C]-5DS whose *m/z* is increased by one mass unit compared with an unlabeled authentic standard in ^13^C-18-HO-MeCLA-fed samples (Fig. 6a). These results suggested that [10-^13^C]-18-HO-MeCLA was converted to [6 -^13^C]-5DS, likely via [10-^13^C]-18-HO-CLA by *L. japonicus*. We assumed that 18-HO-MeCLA is likely to be a precursor also for LL since this non-canonical SL has the B-ring and the carboxylmethyl group. LC-MS/MS analysis of the above exudates revealed that ^13^C-label was incorporated into LL (Fig. 6b). In this case, the bioconversion of [10-^13^C]-18-HO-MeCLA to [6 -^13^C]-LL presumably proceeded without hydrolysis of methyl ester.

**Fig. 6.**
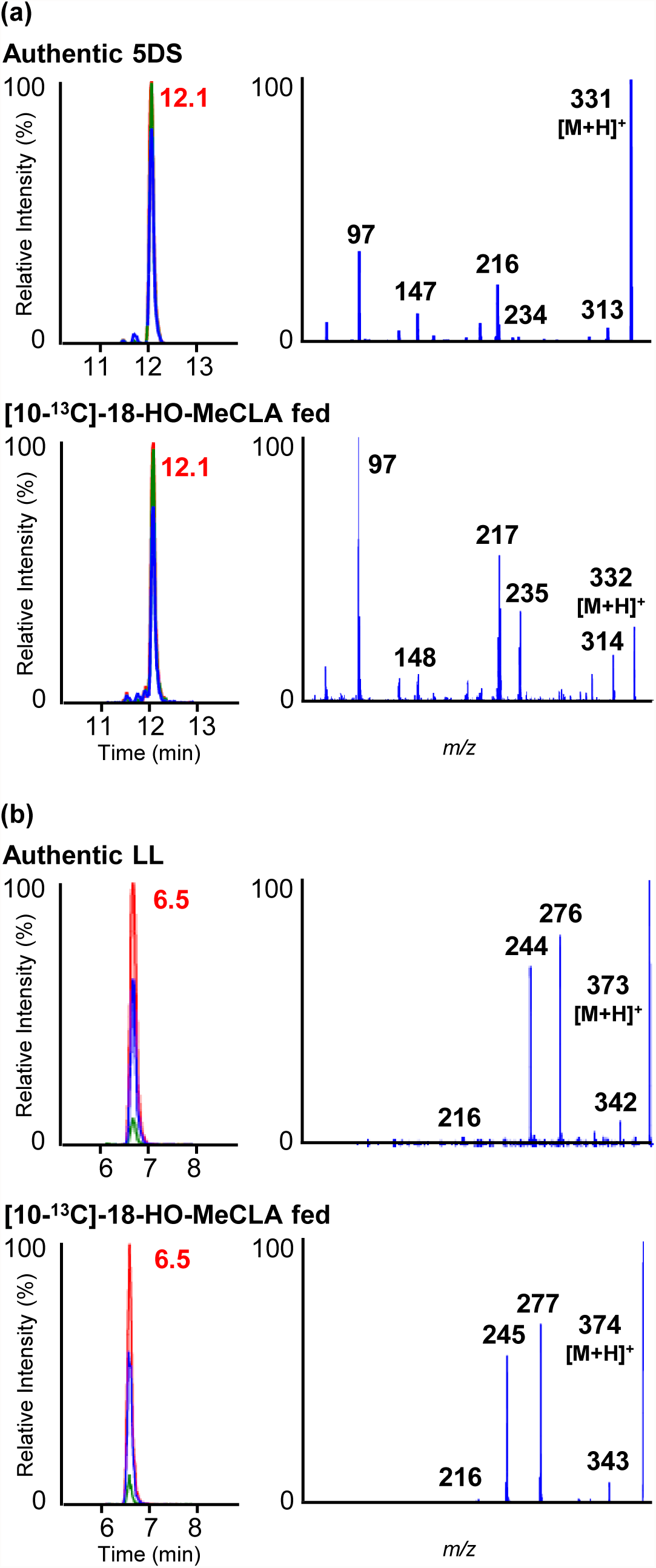
Conversion of [^13^C]-18-HO-MeCLA to [^13^C]-5DS and [^13^C]-lotuslactone (LL) in the feeding experiment using *L. japonicus* roots. (a) Conversion of [10-^13^C]-18-HO-MeCLA to [6 -^13^C]-5DS. LC-MS/MS analysis of [6 -^13^C]-5DS in root exudates after feeding [10-^13^C]-18-HO-MeCLA. MRM chromatograms (left) and full-scan spectra of fragment ions (right). The MRM chromatograms of authentic 5DS (red: 331.15/217.00, blue: 331.15/97.00, green: 331.15/234.00, *m/z* in positive mode) and [6 -^13^C]-5DS (red: 332.15/218.00, blue: 332.15/97.00, green: 332.15/235.00, *m/z* in positive mode) are shown. (b) Conversion of [10-^13^C]-18-HO-MeCLA to [6 -^13^C]-LL. LC-MS/MS analysis of [6 -^13^C]-LL in root exudates after feeding [10-^13^C]-18-HO-MeCLA. MRM chromatograms (left) and full-scan spectra of fragment ions (right). The MRM chromatograms of authentic LL (red: 373.00/276.00, blue: 373.00/244.00, green: 373.00/216.00, *m/z* in positive mode) and [6 -^13^C]-LL (red: 374.00/277.00, blue: 374.00/245.00, green: 374.00/216.00, *m/z* in positive mode) are shown.

## 3. Discussion

The deficiency of 5DS and LL production in the *L. japonicus max1* mutant is similar to those described for mutants in MAX1 homologs in rice, Arabidopsis and tomato (Abe et al., 2014; Cardoso et al., 2014; Zhang et al., 2018). This indicates that LjMAX1 (CYP711A9) is the crucial enzyme for SL biosynthesis in *L. japonicus.* By yeast heterologous expression and chemical synthesis, we demonstrated that LjMAX1 converts CL via CLA to18-HO-CLA but not to 5DS. The inability of LjMAX1 to produce 5DS from CL was confirmed by feeding experiments using recombinant enzymes expressed in microsomes and cells. Therefore, 18-HO-CLA appears to be an end-product by LjMAX1, being classified this enzyme to the A1-type MAX1s such as PtMAX1a and PtMAX1b (Kaori Yoneyama et al., 2018a). In addition to the LjMAX1 (CYP711A9) which we characterized in this study, another putative AtMAX1 homolog (Lj5g3v0408240.1) has been annotated in the recently released *Lotus japonicus* genome assembly build 3.0. The *LjMAX1* gene (chr1.CM0133.560.r2.m) in the build 2.5 was accordingly renamed Lj1g3v1786360.1 in the build 3.0. Future studies are needed to clarify the involvement of the putative homolog in *Lotus* SLs biosynthesis.

The cation-initiated cascade cyclization for constructing the BC ring system of SLs developed by Chojnacka et al. (2011) made a prediction for an intermediate 18-HO-CLA in the enzymatic conversion of CL to 4DO by Os900 (Alder et al. 2012; Zhang et al. 2014; Al-Babili & Bouwmeester, 2015). In our previously study, we detected a new product CLA+16 Da in the reaction of Os900 with CL and CLA (Kaori Yoneyama et al., 2018a). The profile of the product ions of 18-HO-CL metabolite by Os900 was the same as that of CLA+16 Da. These results allowed us to predict that the chemical structure of CLA+16 Da might be 18-HO-CLA. This putative metabolite was produced from CL by SmMAX1a, SmMAX1b, PtMAX1a and PtMAX1b, and also from CLA by the four MAX1s and LjMAX1. In this study, we synthesized 18-HO-MeCLA and compared its chemical properties with CL metabolite by Os900 methylated using diazomethane by LC-MS/MS. Thus, we unambiguously identified CLA+16 Da to be 18-HO-CLA.

In this study, we showed in *L. japonicus* that (11*R*)-CL exists endogenously, and that (11*R*)-CL is stereospecifically converted to (2 *R*)-(+)-5DS. The endogenous presence of (11*R*)-CL has been reported in *Arabidopsis* and rice (Seto et al., 2014). In rice, (11*R*)-CL is converted to (-)-4DO and (-)-orobanchol in a stereospecific manner as in *L. japonicus*. In sorghum, cotton, moonseed and cowpea, the stereospecific conversion of (11*R*)-CL to their respective strigol- and orobanchol-type SLs was demonstrated (Iseki et al., 2018). In tomato, biochemically prepared CL (Alder et al., 2012) was converted to its canonical SLs, orobanchol, *ent*-2 -epiorobanchol and solanacol, and also to the three putative didehydroorobanchol isomers (Zhang et al., 2018).

18-HO-CLA which we identified as the end-product by LjMAX1 is likely to be a precursor for 5DS biosynthesis in *L. japonicus*. This is likely to be the case for the 4DO biosynthesis of popular whose two MAX1 homologs produce 18-HO-CLA as an end-product (Kaori Yoneyama et al., 2018a). To support this hypothesis, we show that 18-HO-CLA is produced from CLA *in planta* by feeding experiments and also that 18-HO-CLA is endogenously present in the root exudates of *L. japonicus* (Fig. S7). Furthermore, we demonstrated that 18-HO-MeCLA is converted to 5DS *in planta* by feeding experiments. Taken together, these data are first to demonstrate that 18-hydroxylated carlactonoates are the intermediates for canonical SL biosynthesis.

LL is a non-canonical SL that contains the AB-ring but lacks the C-ring. This C_20-_skeleton compound has been presumed to be derived from MeCLA or its isomers and their oxygenated derivatives (Yoneyama et al., 2018b). We showed that LL is produced from 18-HO-MeCLA *in planta* by feeding experiments. This indicates that 18-hydroxylated carlactonoates are common intermediates for 5DS and LL in *L. japonicus*. A probable cyclization mechanism for the two different ring systems between 5DS and LL from the common intermediates can be well explained by the hypothesis for the ring formation reaction (Chojnacka et al., 2011). The single concerted mechanism may form the BC-ring of 5DS, whereas the stepwise mechanism is applicable to explain the formation of the B ring in LL.

## 4. Concluding Remarks

In this study, we showed that LjMAX1 (CYP711A9) is the crucial enzyme in the biosynthesis of *Lotus* SLs and that 18-hydroxylated carlactonoates are precursors for SL biosynthesis in *L. japonicus* (Fig. 7). Considering the structure of 18-hydroxylated carlactonoates and the mechanisms of cyclization reaction, the downstream enzyme(s) involved in 5DS biosynthesis might catalyze protonation of the hydroxyl and a stereocontrolled cyclization of B-and β-oriented C ring accompanied by the loss of a water molecule. LL is a highly decorated SL, and multiple enzymes should be included in its biosynthesis from 18-hydroxylated carlactonoates. Our discovery will further contribute to the understanding of the molecular basis for the structural diversity in SLs.

**Fig. 7.**
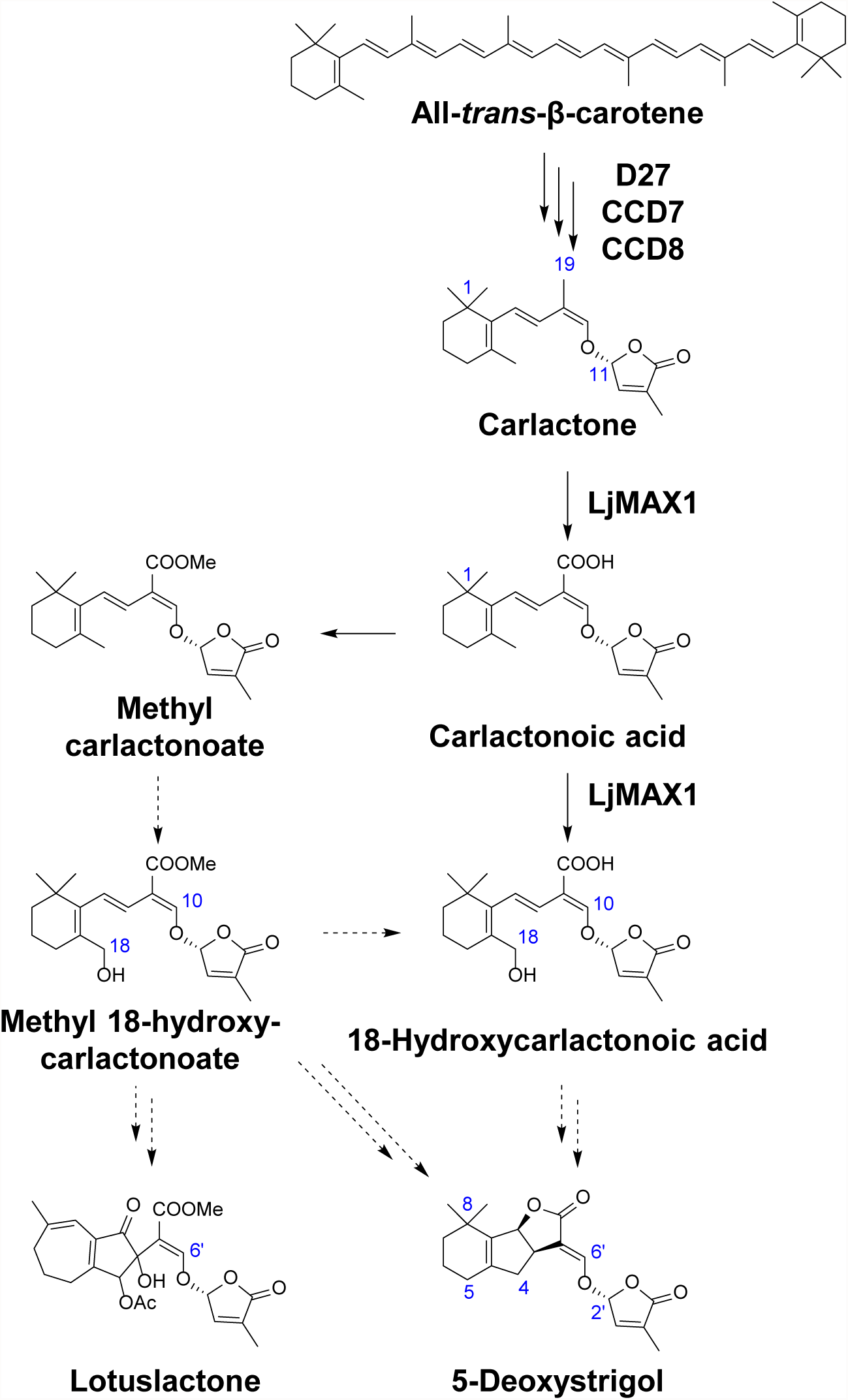
The proposed biosynthetic pathway of 5DS and LL in *L. japonicus*. LjMAX1 catalyzes the oxidation of CL to 18-HO-CLA via CLA. (11*R*)-CL, CLA and 18-hydroxylated carlactonoates are precursors for 5DS and LL in *L. japonicus*. Solid arrows indicate confirmed pathways reported in previous studies and this study and dashed arrows indicate putative pathways. Blue letters indicate the position number.

## 5. Experimental

### 5.1 Plant material and growth conditions

*Lotus japonicus* ecotype Gifu B-129 provided by the National BioResource Project (*L. japonicus* and *Glycine max*; https://www.legumebase.brc.miyazaki-u.ac.jp/index.jsp) was used as the WT. *max1* mutant (line 30101744) was obtained from the *LORE1* collection (Fukai *et al.*, 2012; Urbański *et al.*, 2012).

For LjMAX1 cloning, a feeding experiment of 18-HO-MeCLA and analysis of *LjMAX1 LORE1* insertion mutant, 5-wk-old seedlings were used. Scarified seeds were germinated in vermiculite in a 9-cm pot for one week. The seedlings were grown with M medium (pH 6.8) containing 0.035 mM KH_2_PO_4_ (Akiyama et al., 2005) for three weeks in a growth chamber (MLR-352, Panasonic Healthcare) with a 16–/8–h day/night cycle at 24 °C. The 4-wk-old plants were transferred to a Falcon tube (50 ml), and further grown for an additional one week.

For identification of endogenous CL and feeding experiments of CL and CLA, 9-wk-old seedlings were used. Scarified seeds were sown on vermiculite in a plastic tray with a meshed bottom. The tray was placed in a plastic vessel filled with tap water until the germination of the seeds. The hydroponic culture media were changed to 20 L M medium (pH 6.8) containing 0.035 mM KH_2_PO_4_ and grown in a growth chamber (MLR-352, Panasonic Healthcare) with a 16–/8–h day/night cycle at 24 °C for nine weeks after sowing.

### 5.2. Chemicals

^1^H and ^13^C-NMR spectra were obtained with a JNM-AL400 NMR spectrometer (JEOL) and a JNM-ECZ500R NMR spectrometer (JEOL). Chemical shifts were referenced to tetramethylsilane as an internal standard. High-resolution mass spectra were recorded on a MicroTOF-II mass spectrometer (Bruker). Column chromatography was performed with Kieselgel 60 (Merck), Chromatorex ODS (Fuji Silysia Chemical) and Inertsil SIL-100A (ϕ 10 × 250 mm, 5 μm; GL Sciences). *rac*-CL, *rac*-CLA, (*R*)-[1-^13^CH_3_]-CL, (*S*)-[1-^13^CH_3_]*rac*-CL and [1-^13^CH_3_]*rac*-CLA were synthesized previously (Seto et al., 2014; Abe et al., 2014).

### 5.3. LC-MS/MS Analysis

For detection of SLs in root exudates of WT and *max1* mutant and CL-derivatives metabolites by living yeast, LC-MS/MS analysis was performed with UHPLC (Nexera X2; Shimadzu) and an ion trap instrument (amaZon SL, Bruker) with an electrospray source operating in the smart mode. Ion spray voltage was set at 4,500 V in positive ion mode and −3,500 V in negative ion mode. Nitrogen was used as nebulizer (0.5 bar), the flow of drying gas was 4.0 l mim^−1^ and drying gas temperature of 180 °C. The column oven temperature was maintained at 35 °C. The mobile phase consisted of methanol (solvent A) and water (solvent B), both of which contained 0.1% (vol vol^−1^) acetic acid. HPLC separation was conducted with a linear gradient of 35% A (0 min) to 95% A (20 min) for an ODS column (Kinetex C18 column, ϕ 2.1 × 150 mm, 1.7 μm, Phenomenex) at flow rate of 0.2 ml min^−1^.

For detection of CL-derivatives metabolites by microsome, SLs from feeding experiments and endogenous 18-HO-CLA, LC-MS/MS analysis was performed with UHPLC (Nexera X2; Shimadzu) and a triple quadrupole/linear ion trap instrument (LIT) (QTRAP5500; AB SCIEX) with an electrospray source as reported previously (Abe et al., 2014; Xie et al., 2013).

For identification of endogenous CL, LC-MS/MS analysis was performed with UHPLC (Nexera; Shimadzu) and a quadrupole/time-of-flight tandem mass spectrometer (TripleTOF 5600; AB SCIEX) with an electrospray source as reported previously (Seto et al., 2014).

### 5.4. LjMAX1 cloning

The 5-wk-old seedlings were transferred to hydroponic culture with tap water, and further grown for an additional 3 days. Total RNA was extracted from roots using the RNeasy Plant Mini Kit (Qiagen). The first strand cDNA was synthesized from total RNA using SMARTer RACE cDNA Amplification Kit (Clontech). ORF fragments were amplified by Advantage® 2 Polymerase Mix (Clontech) from the cDNA. The primer pairs used were LjMAX1 forward primer (LjMAX1-F) and reverse primer (LjMAX1-R). Primer sequences listed in Table S1 were used. The ORF fragments were ligated into the cloning vector pTAC1 (DynaExpress) for sequencing. The ORF cDNAs were ligated into a BamHI/SacI site of the pYeDP60 vector (Pompon *et al.*, 1996). The resulting pYeDP60-LjMAX1 plasmid was transformed into *Saccharomyces cerevisiae* strain WAT11 (Pompon et al., 1996) by *S. cerevisiae* Direct Transformation Kit (Wako). *S. cerevisiae* strain WAT11 transformed with the pYeDP60-Os900 plasmid (Kaori Yoneyama et al., 2018a) or empty vector were used positive and negative controls, respectively.

### 5.5. Characterization of an *LjMAX1 LORE1* insertion allele and analysis of SL production

An *LjMAX1 LORE1* insertion allele (line 30101744) and segregating WT were genotyped with the primers indicated in Table S1, together with the *LORE1*-5out primer as described (Fukai et al., 2012; Urbański et al., 2012).

The 5-wk-old *Ljmax1 LORE1* insertion mutants were transferred to a Falcon tube (50 ml) containing 40 ml tap water, and were grown for an additional 2 days. The culture media were passed through Oasis HLB column 1cc, and eluted with 3 ml 90% acetone/water with 0.1% acetic acid. The eluates were concentrated to water, and the residues were extracted with ethyl acetate twice. The ethyl acetate phase was concentrated to dryness under nitrogen gas, dissolved immediately in acetonitrile and subjected to LC-MS/MS analysis using UHPLC (Nexera X2; Shimadzu) and an ion trap instrument (amaZon SL, Bruker) as above.

### 5.6. Gene expression analysis of *LjMAX1 LORE1* insertion mutants

Total RNA was extracted from roots of 5-wk-old seedlings using RNeasy Plant Mini Kit (Qiagen). Total RNA was used to synthesize cDNA using the High Capacity RNA-to-cDNA Kit (Invitrogen). Real-time quantitative PCR (qPCR) was performed with the real-time PCR system (Applied Biosystems7300, Applied Biosystems) using primers as listed in Table S1. Relative expression of transcripts was normalized to the average expression level of *L. japonicus* housekeeping gene (*Ubiquitin 4*) amplified using primers as listed in Table S1. To compare gene expression in WT and *LORE1* insertion mutant (*Ljmax1*), the expression levels were normalized to the expression levels in the WT plants.

### 5.7. Heterologous expression of LjMAX1 in yeast and microsome preparation

Heterologous expression in yeast and microsome preparation from yeast was performed as described previously (Abe et al., 2014; Yoneyama et al., 2018a).

### 5.8. Microsomal conversion of CL and CLA

Each 30 μM of CL and CLA was incubated with the prepared microsomes in the presence of 500 μM NADPH at 30°C for 1 h. The reaction mixture was extracted with 1 ml ethyl acetate twice. The ethyl acetate phase was concentrated to dryness under nitrogen gas, dissolved immediately in acetonitrile and subjected to LC-MS/MS analysis using UHPLC (Nexera X2; Shimadzu) and a triple quadrupole/linear ion trap instrument (LIT) (QTRAP5500; AB SCIEX) as above.

### 5.9. Bioconversion of CL and CLA using living yeast cells

Yeast having pYeDP60-LjMAX1 plasmid was grown in 1 ml YPGE medium at 30°C in a shaking incubator (160 rpm) for 14 h. Galactose was added at the final concentration of 20 g/l and further incubated with shaking at 30°C for 12 h. CL or CLA was added at the final concentration of 30 μM and further incubated with shaking at 30°C for 3 h. The reaction mixture was extracted with 1 ml ethyl acetate twice. The residue was subjected to silica gel column chromatography (Kieselgel 60, Merck) eluted successively with 20% ethyl acetate in *n*-hexane and 100% ethyl acetate with 0.1% acetic acid. The 100% ethyl acetate with 0.1% acetic acid eluate was concentrated to dryness under nitrogen gas, dissolved immediately in acetonitrile and subjected to LC-MS/MS analysis using UHPLC (Nexera X2; Shimadzu) and an ion trap instrument (amaZon SL, Bruker) as above.

### 5.10. Methyl ester derivatization of CL metabolites by Os900

*rac*-CL was incubated with yeast microsomes expressing *Oryza sativa* Os900 (Yoneyama *et al*., 2018a). The extract of the microsomes was treated with ethereal diazomethane for 30 min at room temperature to esterify free carboxylic acids. After evaporation of the solvents under nitrogen gas, the concentrate was dissolved in acetonitrile and subjected to LC-MS/MS analysis using UHPLC (Nexera X2; Shimadzu) and a triple quadrupole/linear ion trap instrument (LIT) (QTRAP5500; AB SCIEX) as above.

### 5.11. Chemical synthesis of ^13^C-labeled and unlabeled 18-HO-MeCLA

#### 5.11.1. 2-(Hydroxymethyl)-6,6-dimethylcyclohex-1-en-1-yl trifluoromethanesulfonate (3)

To a solution of methyl 1,3-dimethyl-2-(((trifluoromethyl)sulfonyl)oxy)cyclohex-2-ene-1-carboxylate **2** (13.7 g, 43.3 mmol) in THF (152 ml) was added a solution of diisobutylaluminium hydride (1.0 M, 126 ml, 126 mmol) in hexane at −78 °C under Ar, and stirred at the same temperature for 30 min. The mixture was allowed to warm to room temperature and stirred for 1.0 h. The mixture was cooled 0 °C and added sat. NH_4_Cl. The suspension was filtrated and extracted with ether. The organic phase was washed with brine, dried with. Na_2_SO_4_ and concentrated in vacuo to give the hydroxy triflate as a blown oil (11.8 g, 40.8 mmol, 94%). ^1^H-NMR (CDCl_3,_ 400 MHz) δ: 1.18 (3H, s, 6-CH_3_), 1.61-1.75 (4H, m, H-4, and −5), 2.36 (2H, t, *J* = 6.1 Hz, H-3), 4.14 (2H, d, *J* = 6.6 Hz, 2-CH_2_); ^13^C-NMR (CDCl_3_, 100 MHz) δ: 26.2, 26.3, 28.2, 35.6, 40.3, 60.1, 117.1, 120.3, 129.4, 150.6; HR-ESI-MS m/z: 311.0548 [M+Na]^+^ (calcd. for C_10_H_15_F_3_NaO_4_S^+^, m/z 311.0535).

#### 5.11.2. 2-(((*tert*-Butyldimethylsilyl)oxy)methyl)-6,6-dimethylcyclohex-1-en-1-yl trifluoromethanesulfonate (4)

A mixture of hydroxy-cyclohexanone the hydroxy triflate (11.8 g, 40.8 mmol), *tert*-butyldimethylsilylchlorosilane (12.3 g, 81.6 mmol), and imidazole (11.2 g, 165 mmol) in DMF (118 ml) was stirred at room temperature for 3 h under Ar. The reaction mixture was taken up in 2% (vol/vol) ether in n-hexane, washed with water, dried over anhydrous Na_2_SO_4_, and concentrated in vauo. Purification by silica gel column chromatography eluted stepwise with *n*-hexane and ether (1% (vol/vol) increments) gave the *tert*-butyldimethylsilyloxy triflate (17.5 g, 43.3 mmol, quant.). ^1^H-NMR (CDCl_3,_ 400 MHz) δ: 0.06 (6H, s, Si-(CH_3_)_2_), 0.89 (9H, s, Si-C(CH_3_)_3_), 1.16 (6H, s, 6-(CH_3_)_2_), 1.60-1.68 (4H, m, H-4 and -5), 2.29 (2H, t, *J* = 6.0 Hz, H-3), 4.27 (2H, s, 2-CH_2_); ^13^C-NMR (CDCl_3_, 100 MHz) δ: −5.6, −5.5, 18.3, 25.7 and 25.9, 26.2 and 26.4, 27.2, 35.6, 40.5, 60.5, 117.1, 120.3, 130.1, 148.7; HR-ESI-MS m/z: 425.1400 [M+Na]^+^ (calcd. for C_16_H_29_F_3_NaO_4_SSi^+^, m/z 425.1400).

#### 5.11.3. (*E*)-3-(2-(((*tert*-Butyldimethylsilyl)oxy)methyl)-6,6-dimethylcyclohex-1-en-1-yl)acrylaldehyde (5)

A mixture of *tert*-butyldimethylsilyloxy triflate (17.5 g), tetrabutylammonium chloride (11.3 g, 40.8 mmol), potassium carbonate (11.6 g, 81.6 mmol), acrylaldehyde (11.2 ml, 168 mmol), triphenylphosphine (879 mg, 3.26 mmol) and palladium (II) acetate (379 mg, 1.63 mmol) in DMF (194 ml) and water (19.4 ml) was stirred at 45 °C for 20 h under Ar. The reaction mixture was poured into a solution of hexane and water (1:1, vol/vol), and extracted with EtOAc twice and ether once. The organic phase was washed with brine and water, dried over anhydrous Na_2_SO_4_, and concentrated in vacuo. Purification by silica gel column chromatography using 5% (vol/vol) stepwise elution with ether and *n*-hexane gave the ionone as a yellow oil (3.12 g, 10.1 mmol, two steps, 25%). ^1^H-NMR (CDCl_3,_ 400 MHz) δ: 0.04 (6H, s, Si-(CH_3_)_2_), 0.89 (9H, s, Si-C(CH_3_)_3_), 1.09 (6H, s, 6’-(CH_3_)_2_), 1.50-1.69 (4H, m, H-4’ and 5’), 2.23 (2H, t, *J* = 6.0 Hz, H-3’), 4.13 (2H, s, 2’-CH_2_), 6.14 (1H, dd, *J* = 16.0, 8.0 Hz, H-2), 7.23 (1H, d, *J* = 16.0 Hz, H-3), 9.57 (1H, d, *J* = 8.0 Hz, H-1); ^13^C-NMR (CDCl_3_, 100 MHz) δ: −5.3 and −5.2, 14.0 and 14.2, 18.3 and 18.6, 25.8 and 26.0, 28.5 and 28.6, 28.7, 34.2, 39.2, 64.0, 133.9 and 134.2, 137.8, 139.4, 152.0 and 152.0, 194.0 and 194.1; HR-ESI-MS m/z: 331.2054 [M+Na]^+^ (calcd. for C_18_H_32_NaO_2_Si^+^, m/z 331.2064).

#### 5.11.4. 2-(*E*)-2-[2-((*tert*-Butyldimethylsiloxy)methyl-6,6-trimetylcyclohex-1-en-1-yl)ethenyl] oxirane (6)

To a solution of trimethylsulfonium iodide (3.0 g, 14.7 mmol) in DMSO (12 ml) was added THF (12 ml) under Ar to yield a finely divided suspension of sulfonium salt. This mixture was then cooled to 0 °C and treated with a solution of dimsyl sodium (4.4 M, 4.1 ml, 18.0 mmol). The resulting gray colored suspension was treated with a solution of aldehyde (3.12 g, 10.1 mmol) in THF (1.6 ml). After stirring at 0 °C for 1.0 h, the mixture was warmed to room temperature and stirred for 15 min. The reaction mixture quenched with water and extracted with *n*-hexane. The organic phase was washed with water, dried over anhydrous Na_2_SO_4_, and concentrated in vacuo to give the crude epoxide (3.3 g, quant.) which was used for the next reaction without purification. ^1^H-NMR (CDCl_3,_ 400 MHz) δ: 0.03 (6H, s, Si-(CH_3_)_2_), 0.88 (9H, s, Si-C(CH_3_)_3_), 1.01 (6H, s, 6’’-(CH_3_)_2_), 1.44-1.61 (4H, m, H-4’’- and -5’’), 2.13 (2H, t, *J* = 6.1 Hz, H-3’’), 2.66 (1H, dd, *J* = 5.4, 2.8 Hz, H-1a), 2.99 (1H, dd, *J* = 5.4, 4.1 Hz, H-1b), 3.38-3.41 (1H, m, H-2), 4.13 (2H, d, *J* = 2.8 Hz, CH_2_O), 5.16 (1H, dd, *J* = 15.7, 8.0 Hz, H-2’), 6.34 (1H, d, *J* = 15.7 Hz, H-1’); ^13^C-NMR (CDCl_3_, 100 MHz) δ: −5.2, 18.4, 18.9, 25.9 and 26.0, 27.7, 28.5 and 28.6, 34.0, 39.1, 48.8, 52.6, 64.5, 130.9, 132.0, 133.3, 139.0; HR-ESI-MS m/z: 345.2211 [M+Na]^+^ (calcd. for C_19_H_34_NaO_2_Si^+^, m/z 345.2220).

#### 5.11.5. (*E*)-4-(2-(((*tert*-Butyldimethylsilyl)oxy)methyl)-6,6-dimethylcyclohex-1-en-1-yl)but-3-enal (7)

To a solution of 2,6-di-*tert*-butyl-4-bromophenol (1.15 g, 4.03 mmol) in CH_2_Cl_2_ (75 ml) was added at room temperature a 1.4 M hexane solution of trimethylaluminium (1.52 ml, 2.13 mmol), and the solution was stirred at room temperature for 1 h under Ar. To a solution of the MABR (2.0 mmol) in CH_2_Cl_2_ was added a solution of crude epoxide (3.3 g, crude) in CH_2_Cl_2_ (6.5 ml) at −78 °C, and the resulting mixture was stirred at −78 °C for 30 min under Ar. The reaction mixture was poured into ice-cold water (90 ml), and extracted with *n*-hexane. The organic phase was washed with saturated aqueous NaHCO_3_ and brine, dried over anhydrous Na_2_SO_4_, and concentrated in vacuo. The residue was subjected to ODS column chromatography eluted stepwise with 70–100% (vol/vol) acetonitrile in water [5% (vol/vol) increments]; 90–95% acetonitrile eluates were combined, evaporated to water in vacuo, extracted with ether, and evaporated to give the *tert*-butyldimethylsilyloxy-aldehyde (1.36 g, 4.21 mmol, two steps 42%). ^1^H-NMR (CDCl_3,_ 400 MHz) δ: 0.03 (6H, s, Si(CH_3_)_2_), 0.88 (9H, s, SiC(CH_3_)_3_), 0.99 (6H, s, 6’-CH_3_), 1.59-1.89 (2H, m, H-4’ and 5’), 2.15 (2H, t, *J* = 6.3 Hz, H-3’), 3.23 (2H, dt, *J* = 7.0, 1.9 Hz, H-2), 4.18 (2H, s, 2’-CH_2_), 5.47 (1H, dd, *J* = 15.9, 7.0 Hz, H-3), 6.03 (1H, d, *J* = 15.9 Hz, H-4), 9.71 (1H, t, *J* = 1.9 Hz, H-1); ^13^C-NMR (CDCl_3_, 100 MHz) δ: −5.2, 18.4 and 19.0, 25.8 and 26.1, 30.0 and 30.1, 34.5, 39.0, 47.7, 64.6, 112.6, 127.8 and 127.9, 138.1, 152.9, 199.7; HR-ESI-MS m/z: 345.2223 [M+Na]^+^ (calcd. for C_19_H_34_NaO_2_Si^+^, m/z 345.2220).

#### 5.11.6. Methyl (*E*)-4-(2-(((*tert*-butyldimethylsilyl)oxy)methyl)-6,6-dimethylcyclohex-1-en-1-yl)but-3-enoate (8)

A mixture of aldehyde (1.36 g, 4.21 mmol), sodium cyanide (1.09 g, 22.3 mmol), acetic acid (385 μl, 6.74 mmol), and manganese (IV) oxide (7.32 g, 8.42 mmol) in methanol (67 ml) was stirred at room temperature for 21 h under Ar. The reaction mixture was filtered and evaporated. The residue was partitioned between ether and water. The organic layer was washed with water, dried over MgSO_4_, and concentrated in vacuo. Purification by silica gel column chromatography eluted with *n*-hexane and 1% ether in *n*-hexane gave the carboxylic acid methyl ester (220 mg, 0.623 mmol, 15%). ^1^H-NMR (CDCl_3,_ 400 MHz) δ: 0.02 (6H, s, Si(CH_3_)_2_), 0.88 (9H, s, SiC(CH_3_)_3_), 0.98 (6H, s, 6’-(CH_3_)_2_), 1.43-1.61 (3H, tt, *J* = 9.3, 3.2 Hz), 2.13 (2H, dd, *J* = 6.2, 4.5 Hz), 3.13 (2H, dd, *J* = 7.1, 1.3 Hz), 3.69 (3H, s), 4.19 (2H, s, 2-CH_2_), 5.45 (1H, dt, *J* = 15.6, 7.1 Hz), 5.98 (1H, d, *J* = 15.6 Hz); ^13^C-NMR (CDCl_3_, 100 MHz) δ: −5.3 and −5.2, 18.4 and 19.0, 25.9 and 26.0, 27.3, 28.4 and 28.6, 34.0, 38.3, 39.0, 51.7 and 51.9, 64.5, 125.16 and 125.19, 125.5, 126.0 and 126.3, 130.3, 130.8 and 130.9, 132.8, 139.1, 172.2; HR-ESI-MS m/z: 375.2330 [M+Na]^+^ (calcd. for C_20_H_36_NaO_3_Si^+^, m/z 375.2326).

#### 5.11.7. Methyl (2*E*,3*E*)-4-(2-(((*tert*-butyldimethylsilyl)oxy)methyl)-6,6-dimethylcyclohex-1-en-1-yl)-2-(((4-methyl-5-oxo-2,5-dihydrofuran-2-yl)oxy)methylene)but-3-enoate (18-TBSO-MeCLA, 9a)

To a suspension of sodium hydride (22.5 mg, 0.562 mmol) in DMF (335 μl) was added a solution of carboxylic acid methyl ester (110 mg, 0.312 mmol) in DMF (335 μl) at room temperature under Ar. Then methyl formate (96.5 μl, 1.56 mmol) was added, and the mixture was stirred for 13.5 h. After cooling to 0 °C, (±)-4-bromo-2-methyl-2-buten-4-olide (36.5 μl, 374 mmol) in DMF (84 μl) was added, and the reaction mixture was stirred at room temperature for 3 h under Ar. The mixture was poured into ice cooled 0.1 N HCl and extracted with ether. The organic phase was washed with water, dried over anhydrous Na_2_SO_4_, and concentrated in vacuo. The residue was subjected to silica gel chromatography eluted stepwise with *n*-hexane and ethyl acetate [10% (vol/vol) increments]. The 20-30% ethyl acetate eluates containing crude 18-TBSO-MeCLA were purified by a semipreparative Inertsil SIL-100A HPLC column (ϕ 10 × 250 mm, 5 μm; GL Sciences), using isocratic elution with 5% ethanol in *n*-hexane at a flow rate of 4.0 ml/min and monitored at 254 nm to give 18-TBSO-MeCLA (3.02 mg, 6.34 μmol, 2.0%, ^*t*^R 10.7 min). ^1^H-NMR (C_6_D_6,_ 400 MHz) δ: 0.08 (6H, s, Si(CH_3_)_2_), 0.97 (9H, s, SiC(CH_3_)_3_), 1.09 (3H, s, CH_3_-16 or 17), 1.10 (3H, s, CH_3_-16 or 17), 1.35 (3H, t, *J* = 1.5 Hz, CH_3_-15), 1.40–1.60 (4H, m, H-2 and -3), 2.22-2.37 (2H, m, H-4), 3.40 (3H, s, COOMe), 4.34 (1H, d, *J* = 11.1 Hz, 18-CHa), 4.42 (1H, d, *J* = 11.1 Hz, 18-CHb), 5.05 (1H, m, H-11), 5.73 (1H, m, H-12), 6.54 (1H, d, *J* = 16.3 Hz, H-8), 7.26 (1H, br. d, *J* = 16.3 Hz, H-7), 7.46 (1H, s, H-10); ^13^C-NMR (C_6_D_6_, 100 MHz) δ: −5.08, −5.03, 10.2, 18.6, 19.4, 19.9, 26.2, 28.2, 28.9, 34.5, 39.5, 51.2, 65.2, 100.4, 112.8, 122.9, 128.52, 131.9, 132.9, 135.11, 140.6, 141.5, 152.5, 166.5, 169.7, 178.7; HR-ESI-MS m/z: 499.2461 [M+Na]^+^ (calcd. for C_26_H_40_NaO_6_Si^+^, m/z 499.2486).

#### 5.11.8. Methyl (2*E*,3*E*)-4-(2-(hydroxymethyl)-6,6-dimethylcyclohex-1-en-1-yl)-2-(((4-methyl-5-oxo-2,5-dihydrofuran-2-yl)oxy)methylene)but-3-enoate (18-HO-MeCLA, 1a)

To a solution of 18-TBSO-MeCLA (6.0 mg, 13.0 μmol) in THF (1.5 ml) was added at room temperature acetic acid (4.5 ml) and water (1.5 ml), and the mixture was stirred for 3 h. The reaction mixture was poured into water and extracted with ether. The organic phase was washed with water, dried over anhydrous Na_2_SO_4_, and concentrated in vacuo. Purification by silica gel column chromatography using 20% (vol/vol) stepwise elution with ether and *n*-hexane gave the 18-HO-MeCLA as a yellow oil [3.1 mg, 8.6 μmol, 66%]. 18-HO-MeCLA ^1^H-NMR (C_6_D_6,_ 400 MHz) δ: 1.07 (3H, s, CH_3_-16 or 17), 1.10 (3H, s, CH_3_-16 or 17), 1.31 (3H, t, *J* = 1.5 Hz, CH_3_-15), 1.37–1.56 (4H, m, H-2 and -3), 2.11-2.14 (2H, m, H-2), 3.39 (3H, s, COOMe), 4.11 (1H, d, *J* = 11.5 Hz, 18-CHa), 4.16 (1H, d, *J* = 11.5 Hz, 18-CHb), 5.05 (1H, m, H-11), 5.70 (1H, m, H-12), 6.51 (1H, d, *J* = 16.5 Hz, H-8), 7.24 (1H, br. d, *J* = 16.5 Hz, H-7), 7.46 (1H, s, H-10); ^13^C-NMR (C_6_D_6_, 100 MHz) δ:10.2, 19.4, 19.9, 28.4, 28.9, 34.5, 39.4, 51.2, 64.6, 100.3, 122.8, 132.0, 133.1, 135.1, 140.7, 142.5, 152.5, 166.5, 169.8, 175.8; HR-ESI-MS m/z: 385.1634 [M+Na]^+^ (calcd. for C_20_H_26_NaO_6_ ^+^, m/z 385.1622).

#### 5.11.9. Methyl (2*E*,3*E*)-4-(2-(((*tert*-butyldimethylsilyl)oxy)methyl)-6,6-dimethylcyclohex-1-en-1-yl)-2-(((4-(methyl-^13^C)-5-oxo-2,5-dihydrofuran-2-yl)oxy)methylene)but-3-enoate ([10-^13^C]-18TBSO-MeCLA, 9b)

To a suspension of sodium hydride (21.8 mg, 0.54 mmol) in DMF (294 μl) was added a solution of carboxylic acid methyl ester (95.8 mg, 0.272 mmol) in DMF (478 μl) at room temperature under Ar. Then ^13^C-methyl formate (85.5 μl, 1.36 mmol) was added, and the mixture was stirred for 3 h. After cooling to 0 °C, (±)-4-bromo-2-methyl-2-buten-4-olide (32.1 μl, 0.326 mmol) in DMF (73.5 μl) was added, and the reaction mixture was stirred at room temperature for 1 h under Ar. The mixture was poured into ice cooled 0.2 N HCl and extracted with ether. The organic phase was washed with water, dried over anhydrous Na_2_SO_4_, and concentrated in vacuo. The residue was subjected to silica gel chromatography eluted stepwise with *n*-hexane and ethyl acetate [4% (vol/vol) increments]. The 16-20% ethyl acetate eluates containing crude ^13^C-18-TBSO-MeCLA were purified by a semipreparative Inertsil SIL-100A HPLC column (ϕ 10 × 250 mm, 5 μm; GL Sciences), using isocratic elution with 4% ethanol in *n*-hexane at a flow rate of 4.0 ml/min and monitored at 254 nm to give 18-TBSO-MeCLA (3.86 mg, 8.08 μmol, 2.97%, ^*t*^R 10.2 min). ^1^H-NMR (C_6_D_6,_ 400 MHz) δ: 0.08 (6H, s, Si(CH_3_)_2_), 0.98 (9H, s, SiC(CH_3_)_3_), 1.09 (3H, s, CH_3_-16 or 17), 1.10 (3H, s, CH_3_-16 or 17), 1.34 (3H, t, *J* = 1.5 Hz, CH_3_-15), 1.40–2.31 (6H, m, H-2, -3 and -4), 3.39 (3H, s, COOMe), 4.35 (1H, d, *J* = 11.2 Hz, 18-CHa), 4.42 (1H, d, *J* = 11.2 Hz, 18-CHb), 5.03 (1H, m, H-11), 5.71 (1H, m, H-12), 6.54 (1H, d, *J* = 16.0 Hz, H-8), 7.27 (1H, br. d, *J* = 16.0 Hz, H-7), 7.47 (1H, s, *J* = 187.0 Hz, H-10); ^13^C-NMR (C_6_D_6_, 100 MHz) δ: −5.09, −5.04, 10.2, 18.6, 19.4, 19.9, 26.2, 28.2, 28.9, 34.5, 39.5, 51.2, 65.2, 100.4, 122.9, 131.9, 132.9, 135.1, 140.6, 141.5, 152.5, 166.6, 169.7, 178.7; HR-ESI-MS m/z: 500.2521 [M+Na]^+^ (calcd. for C_25_^13^CH_40_NaO_6_Si^+^, m/z 500.2520).

#### 5.11.10. Methyl (2*E*,3*E*)-4-(2-(hydroxymethyl)-6,6-dimethylcyclohex-1-en-1-yl)-2-(((4-(methyl-^13^C)-5-oxo-2,5-dihydrofuran-2-yl)oxy)methylene)but-3-enoate ([10-^13^C]-18-HO-MeCLA, 1b)

To a solution of ^13^C-18-TBSO-MeCLA (3.36 mg, 7.03 μmol) in THF (0.81 ml) was added at room temperature acetic acid (2.4 ml) and water (0.81 ml), and the mixture was stirred for 3 h. The reaction mixture was poured into water and extracted with ether. The organic phase was washed with water, dried over anhydrous Na_2_SO_4_, and concentrated in vacuo. Purification by silica gel column chromatography using 20% (vol/vol) stepwise elution with EtOAc and n-hexane gave the 18-HO-MeCLA as a yellow oil [1.01 mg, 2.78 μmol, 40%]. [10-^13^C]-18-HO-MeCLA ^1^H-NMR (C_6_D_6,_ 400 MHz) δ: 1.09 (3H, s, CH_3_-16 or 17), 1.12 (3H, s, CH_3_-16 or 17), 1.30 (3H, t, *J* = 1.7 Hz, CH_3_-15), 1.35–1.55 (4H, m, H-2 and -3), 2.13-2.15 (2H, m, H-4), 3.40 (3H, s, COOMe), 4.12 (1H, d, *J* = 11.5 Hz, 18-CHa), 4.17 (1H, d, *J* = 11.5 Hz, 18-CHb), 5.02-50.3 (1H, m, H-11), 5.67 (1H, s, H-12), 6.54 (1H, dd, *J* = 16.6, 4.6, Hz, H-8), 7.27 (1H, br. d, *J* = 16.6 Hz, H-7), 7.47 (1H, d, *J* = 187.4 Hz, H-10); HR-ESI-MS m/z: 386.1657 [M+Na]^+^ (calcd. for C_19_^13^CH_26_NaO_6_, m/z 386.1655).

### 5.12. Identification of endogenous CL

Roots of 9-wk-old seedlings (1–1.5 g) were used. The CL purification method and LC-MS/MS analysis was performed as described previously (Seto et al., 2014).

### 5.13. Identification of 18-HO-CLA in root exudate

The 5-wk-old plants were transferred to a Falcon tube (50 ml) containing 40 ml tap water, and grown for an additional 2 days. The culture media were pass through Oasis HLB column 1cc (Waters), and eluted with 3 ml 90% acetone/water with 0.1% acetic acid. The eluates were concentrated to water, and the aqueous residues were extracted with ethyl acetate twice. The ethyl acetate phase was concentrated to dryness under nitrogen gas, dissolved immediately in acetonitrile, and subjected to LC-MS/MS analysis using UHPLC (Nexera X2; Shimadzu) and a triple quadrupole/linear ion trap instrument (LIT) (QTRAP5500; AB SCIEX) as above.

### 5.14. Feeding of CL and CLA to *L. japonicus* plant

The 9-wk-old plants (*n* = 3) were transferred to glass pots containing Pi-free M medium (100 ml) containing 1 μM (*R*)- or (*S*)-[1-^13^CH_3_]-CL or [1-^13^CH_3_]-CLA and 10 μM fluridone and 0.02% acetone and were grown for an additional 3 days. Control plants were grown in the same volume of hydroponic culture media containing 0.02% acetone. The hydroponic solutions were passed through an activated charcoal column and eluted with acetone. The eluates were concentrated to water in vacuo, and solid K_2_HPO_4_ was added to the aqueous residues at the final concentration of 0.2 M. The aqueous residues were extracted with ethyl acetate. The ethyl acetate phase was concentrated to dryness under nitrogen gas. The concentrates were subjected to silica gel column chromatography (Kieselgel 60, Merck) eluted successively with 20% and 40% ethyl acetate in *n*-hexane, 100% ethyl acetate with 0.1% acetic acid. The fractions eluted with 40% and 100% ethyl acetate were concentrated to dryness under nitrogen gas, dissolved immediately in acetonitrile and subjected to LC-MS/MS analysis using UHPLC (Nexera X2; Shimadzu) and a triple quadrupole/linear ion trap instrument (LIT) (QTRAP5500; AB SCIEX) as above.

### 5.15. Feeding of 18-HO-MeCLA to *L. japonicus* plant

The 5-wk-old plants were transferred to a Falcon tube (50 ml) containing 40 ml tap water with 1 μM [10-^13^C]-18-HO-MeCLA and 10 μM fluridone and 0.02% acetone and were grown for an additional 2 days. Control plants were grown in the same volume of tap water containing 0.02% acetone. The culture media were passed through Oasis HLB column 1cc, and eluted with 3 ml 90% acetone/water with 0.1% acetic acid. The eluates were concentrated to water, and the residues were extracted with ethyl acetate twice. The ethyl acetate phase was concentrated to dryness under nitrogen gas, dissolved immediately in acetonitrile and subjected to LC-MS/MS analysis using UHPLC (Nexera X2; Shimadzu) and a triple quadrupole/linear ion trap instrument (LIT) (QTRAP5500; AB SCIEX) as above.

### 5.16. Accession number

The accession number for the *LjMAX1* sequence used in this study is LC495384.

## Acknowledgments

We thank Denis Pompon (CNRS) for providing pYeDP60 vector and yeast strain WAT11. This work was supported by the Program for Promotion of Basic and Applied Research for Innovations in Bio-Oriented Industry, the Japan Science and Technology Research Promotion Program for Agriculture, Forestry, Fisheries and Food Industry, and Grant-in-Aid for JSPS Fellows [Grant Number 17J05519].

---

**Scheme 1.**
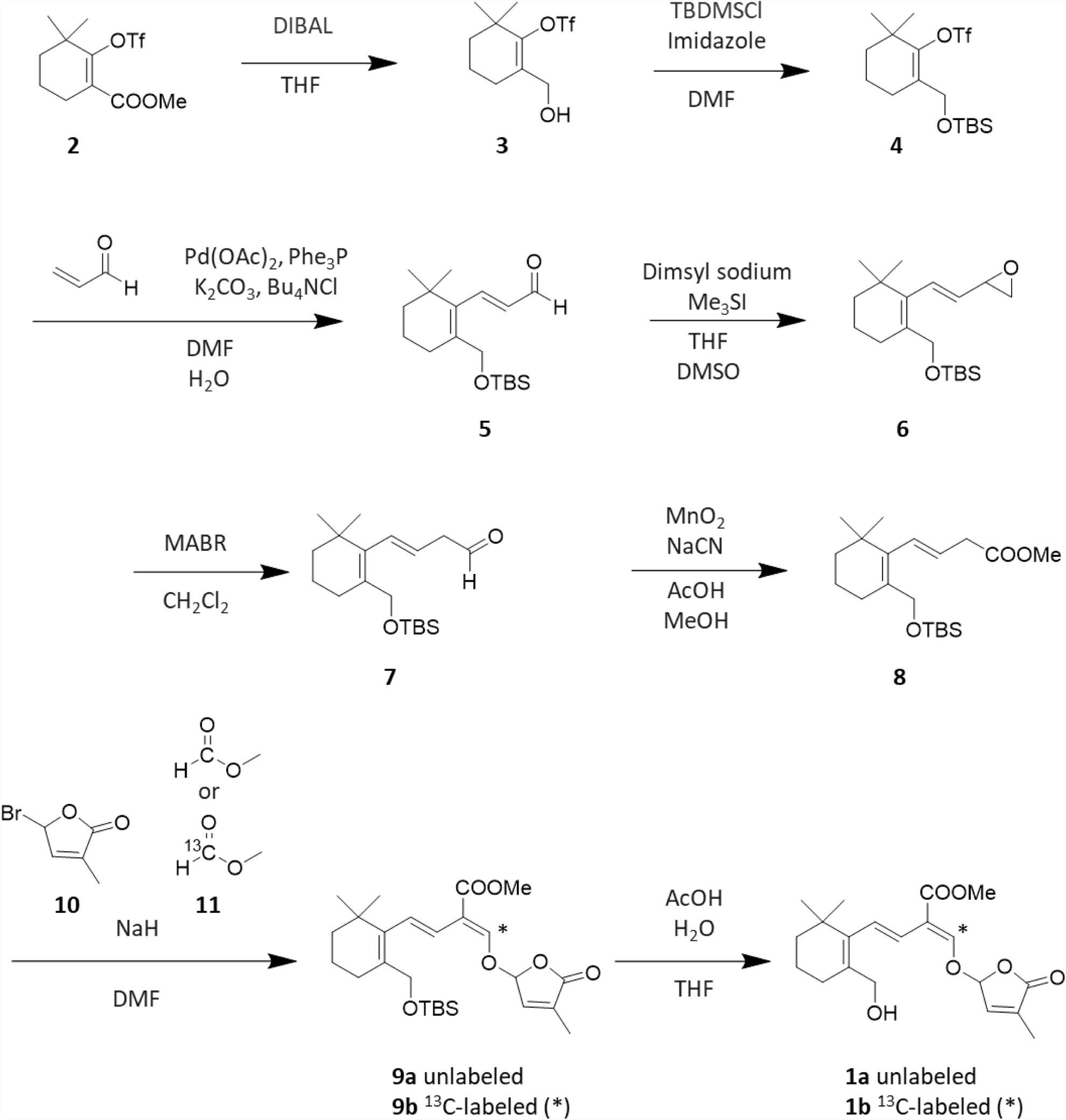
Synthetic scheme of ^13^C-labeled and unlabeled 18-HO-MeCLA. Asterisks indicate the position of ^13^C.

